## Supplementary Figure for "Supramolecular Biomimetic Topology: Macrophage-Engaged Neoadjuvant Platform for Preoperative Immunotherapy"

**Biomimetic Neoadjuvant Platform: Controlled Nanotoxicology as a Novel Approach for Preoperative Immunomodulation**

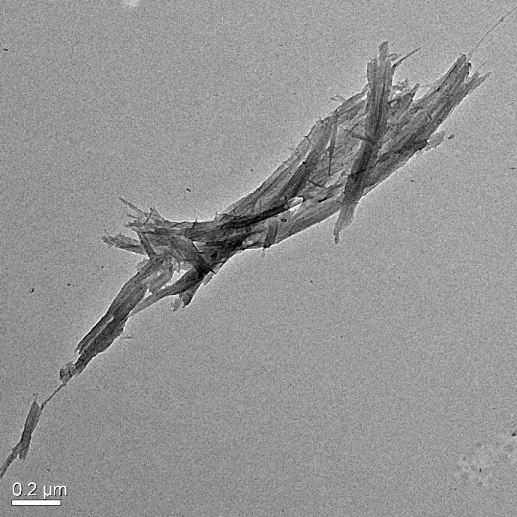

Supplementary Fig. 1. TEM images of non-cut SWNTs exhibited pronounced aggregation

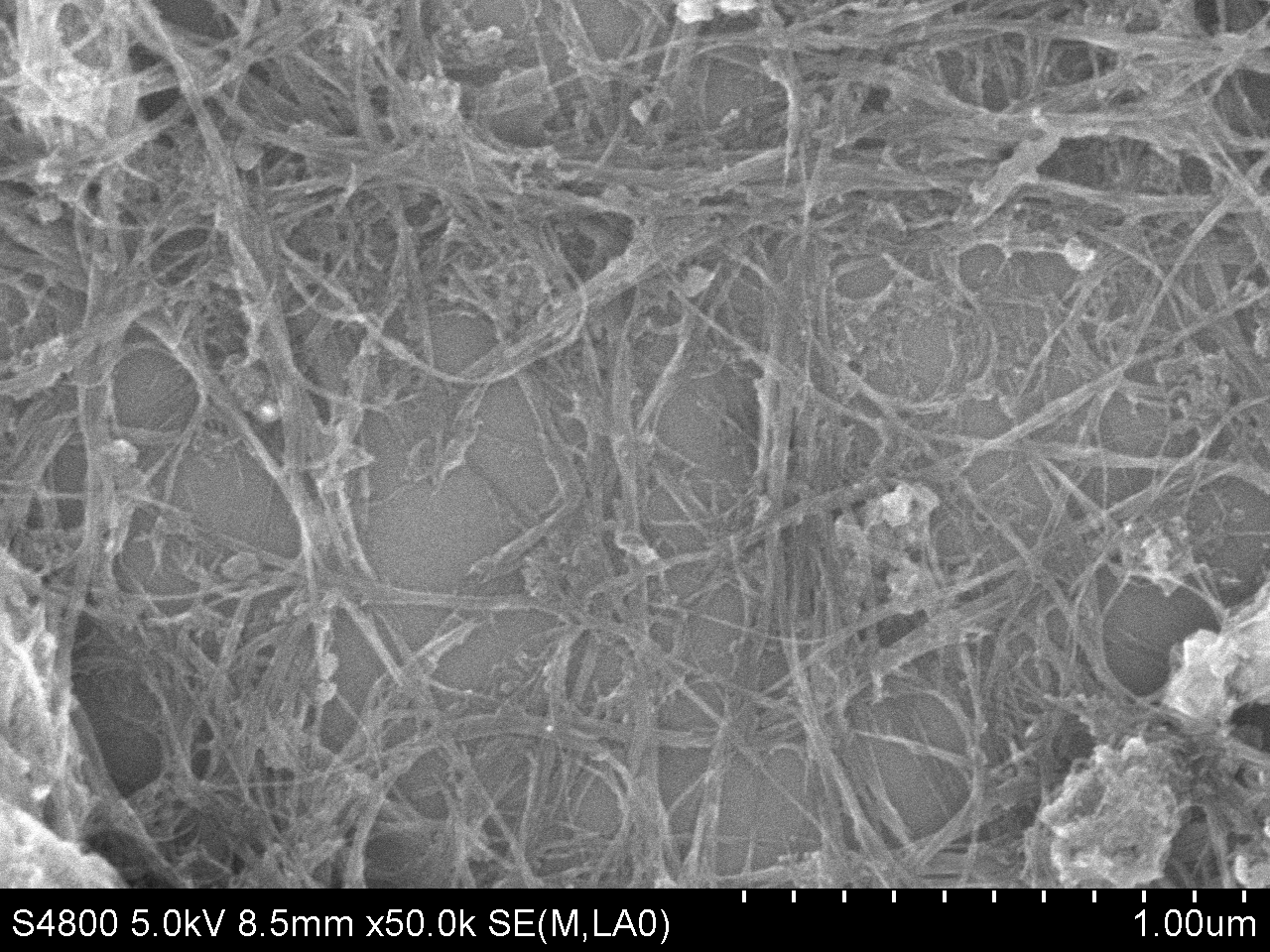

Supplementary Fig. 2. SEM images of non-cut SWNTs exhibited pronounced aggregation

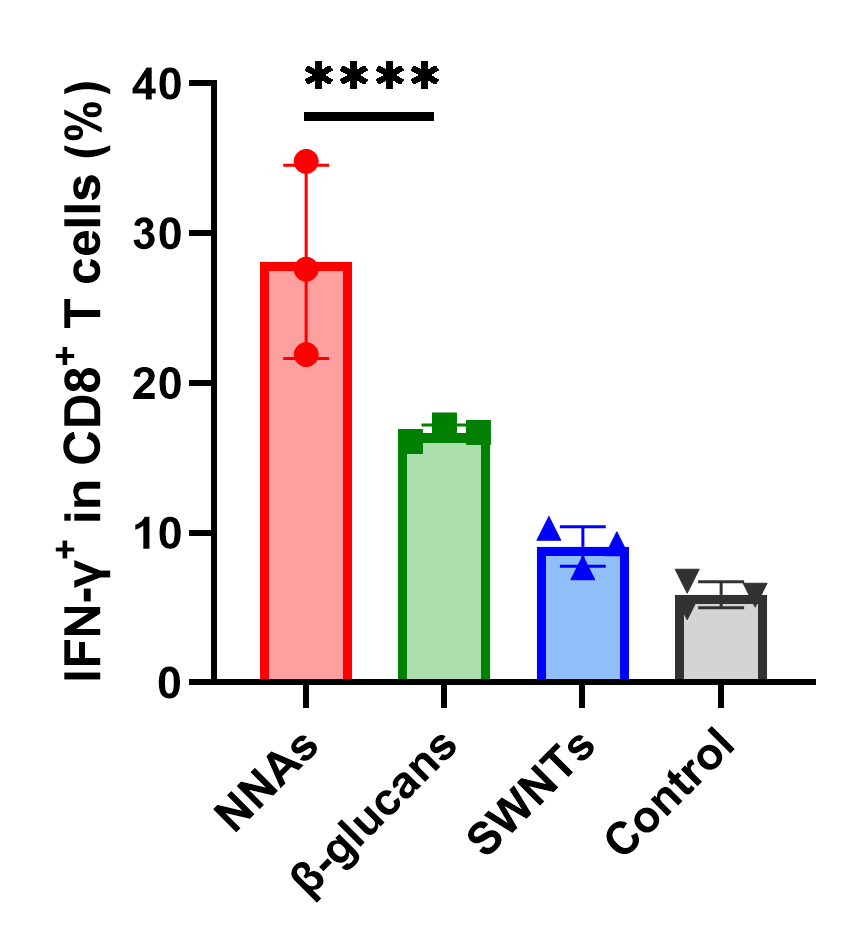

Supplementary Fig. 3. IFN-γ^+^ CD8^+^ T cells
